# Bidirectional acetate exchange couples ethanol turnover to acetyl-CoA metabolism in *Saccharomyces cerevisiae*

**DOI:** 10.64898/2026.11.13.766108

**Authors:** Anthony Carneiro, Mohammad Zishaan Kodooruth, Hanna Kulyk, Gilles Truan, Hélène Martin-Yken, Pierre Millard, Sara Castaño-Cerezo

**Affiliations:** TBI, Université de Toulouse, CNRS, INRAE, INSA, Toulouse, France; MetaboHUB-MetaToul, National Infrastructure of Metabolomics and Fluxomics, Toulouse, France

## Abstract

Overflow metabolism describes the excretion of partially oxidized metabolites into the extracellular environment during rapid sugar utilization. Although these metabolites can later be reused, it remains unclear whether reassimilation also occurs during ongoing sugar consumption, and if so, how. Here, ^13^C-isotope tracing and metabolic flux analysis of acetate exchange in *Saccharomyces cerevisiae* during hexose utilization reveal that net acetate overflow is only the small difference between order-of-magnitude larger simultaneous excretion and assimilation fluxes. Unlike ethanol production, acetate exchange is not determined by hexose uptake but depends on metabolic processes downstream of glycolysis. Ethanol oxidation contributes to acetate excretion even during net ethanol production, directly linking the two main overflow pathways. We exploited this understanding to redirect extracellular acetate into the acetyl-CoA-derived product mevalonate. These results establish acetate overflow as a dynamic exchange linking ethanol turnover to acetyl-CoA metabolism, in which acetate is simultaneously an overflow product and a carbon source that can be redirected toward bioproduction.

## Introduction

Overflow metabolism, the excretion of incompletely oxidized metabolites, is widespread across bacteria, yeasts and mammalian cells. Canonical examples include acetate in *Escherichia coli*^1^, ethanol in *Saccharomyces cerevisiae*^2^ and lactate in mammalian cells^3^. Overflow metabolites have traditionally been considered waste by-products, often with inhibitory or toxic effects^4–6^: excreted while preferred sugars are abundant, but reassimilated, if at all, only once sugars have been depleted^7–9^. This view is now changing. ^13^C-isotope-tracing studies have revealed substantial assimilation of overflow metabolites even as extracellular concentrations increase^10–13^. Moreover, their physiological roles extend beyond carbon disposal^14^. Lactate can serve as an oxidative fuel in mammalian tissues^15^, extracellular ethanol contributes substantially to acetyl-CoA metabolism and redox homeostasis in yeast^11^, and acetate can enhance growth and metabolic adaptation in *E. coli*^7,16^. Rather than mere glycolytic waste, overflow metabolites are therefore emerging as recyclable nutrients that influence cellular physiology.

In *S. cerevisiae*, extracellular ethanol is rapidly reassimilated and contributes to carbon and redox metabolism even during fermentative growth and net ethanol accumulation^11^. *S. cerevisiae* also excretes acetate during growth on glucose^17^. Under these conditions, acetate production is conventionally attributed to the pyruvate dehydrogenase bypass, in which pyruvate-derived acetaldehyde is oxidized to acetate by acetaldehyde dehydrogenase (Ald)^17^, and acetyl-CoA synthetase (Acs) converts acetate into acetyl-CoA^18^ (Fig. 1A). Acetate is thought to be excreted while glucose remains available and reassimilated only after glucose depletion, consistent with glucose-induced catabolite repression of pathways required for acetate utilization^18,19^. However, acetate can cross the plasma membrane in both directions, by passive diffusion and via transporters^20^, and extracellular measurements only report net fluxes. The case of ethanol therefore raises the possibility that acetate is also reassimilated during glucose utilization.

**Fig. 1.**
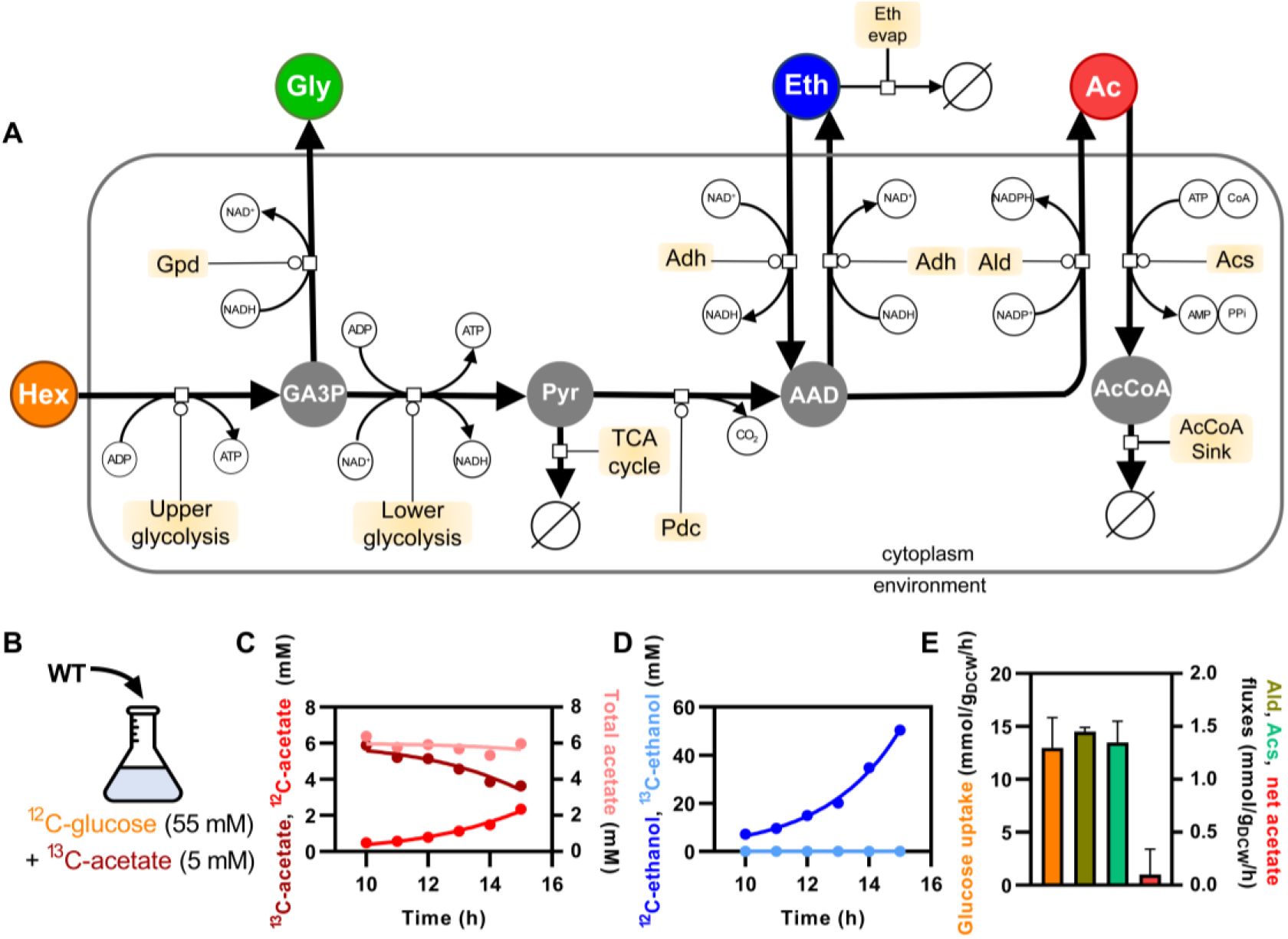
Net acetate overflow masks large opposing fluxes. **(A)** Network of the isotopic model of *S. cerevisiae* central carbon metabolism used for flux estimation, in Systems Biology Graphical Notation (http://sbgn.org). **(B)** Experimental design: cells grown on 55 mM glucose supplemented with 5 mM [U-^13^C]acetate. **(C, D)** Concentration time courses of **(C)** ^12^C-acetate, ^13^C-acetate, and total acetate, and **(D)** ^12^C-ethanol and ^13^C-ethanol during exponential growth. Dots represent one replicate and lines the best fit of the model. **(E)** Glucose uptake, Ald, Acs and net acetate fluxes during exponential growth. Error bars represent [SD; n = 3 biological replicates]. Hex, hexose; GA3P, glyceraldehyde-3-phosphate; Gly, glycerol; Pyr, pyruvate; Eth, ethanol; AAD, acetaldehyde; Ac, acetate; AcCoA, acetyl-CoA; Gpd, glycerol-3-phosphate dehydrogenase; Pdc, pyruvate decarboxylase; Adh, alcohol dehydrogenase; Ald, acetaldehyde dehydrogenase; Acs, acetyl-CoA synthetase; TCA, tricarboxylic acid; Eth evap, ethanol evaporation; ∅, sink.

A few observations suggest that acetate production and utilization are not always sequential. In yeast, glucose and acetate co-utilization has been reported in carbon-limited chemostats, where low residual glucose relieves glucose repression^21^. Furthermore, engineered yeast can efficiently utilize acetate together with xylose^19^. Reassimilated ethanol could also be oxidized through acetaldehyde to acetate, providing an alternative source of acetate during fermentative metabolism. However, these observations under specific metabolic conditions or from engineered strains leave a central question unanswered: what determines the direction of net acetate exchange during sugar utilization?

Net acetate flux, the quantity conventionally measured, cannot answer these questions, because it does not distinguish one-way production from simultaneous production and assimilation. In *E. coli*, acetate production and uptake largely proceed through the same reversible Pta-AckA pathway, and net acetate flux responds strongly to extracellular acetate concentration and glycolytic flux^7,10^. In yeast, however, acetate production and assimilation involve distinct enzymatic routes^17,18^. How this different pathway organization shapes acetate exchange, and whether the principles established in *E. coli* therefore apply to yeast, remain unclear.

Here, we combine ^13^C-isotope tracing and metabolic flux analysis with genetic, chemical and environmental perturbations to resolve acetate exchange in *S. cerevisiae*. We show that net acetate overflow hides a bidirectional exchange, identify what controls the balance between production and assimilation, and demonstrate that ethanol oxidation contributes substantially to acetate production, directly linking the two main overflow pathways of yeast. Finally, we use this knowledge to channel acetate into mevalonate biosynthesis.

## Results

### Net acetate overflow masks large opposing fluxes

During exponential aerobic growth on glucose, CEN.PK2-1D *S. cerevisiae* cells produced ethanol, glycerol, and acetate, with yields of 0.55 ± 0.02, 0.02 ± 0.01, and 0.03 ± 0.01 (C-mol/C-mol), respectively. Ethanol accumulated from the onset of glucose consumption, whereas extracellular glycerol and acetate production began only after a 3 h delay. Extracellular ethanol and acetate were then re-consumed only after glucose depletion (Supplementary Fig. 1). Judged by extracellular concentrations alone, acetate therefore appears to accumulate during glucose growth and be reassimilated after glucose exhaustion.

To investigate whether net acetate excretion masks simultaneous acetate assimilation, we supplemented glucose cultures with 5 mM [U-^13^C]acetate, roughly the maximal extracellular acetate concentration reached with glucose as sole carbon source (Fig. 1B; Supplementary Fig. 1). Total extracellular acetate concentration remained nearly constant (5.8 ± 0.3 mM; Fig. 1C), but unlabeled acetate derived from glucose accumulated as the initial ^13^C-acetate pool decreased, indicating that cells assimilated extracellular acetate while simultaneously producing acetate from glucose. No ^13^C-ethanol was detected, so acetate was not measurably converted back to ethanol under these conditions (Fig. 1D).

To quantify these opposing processes, we used an isotopic model of *S. cerevisiae* central carbon metabolism to separately estimate acetate production through acetaldehyde dehydrogenase (Ald) and acetate consumption through acetyl-CoA synthetase (Acs; Fig. 1A). Production and consumption fluxes were respectively 15- and 14-fold larger than the net acetate flux, and accounted for 11% and 10% of the glucose uptake flux (Fig. 1E). Despite modest net accumulation of acetate in the medium, acetate exchange therefore represents a substantial carbon flux: net overflow is only the difference between two much larger opposing fluxes, reflecting extensive turnover rather than one-way secretion.

### Acetate exchange is uncoupled from glycolytic flux and shifts toward assimilation with respiratory metabolism

In *E. coli*, extracellular acetate concentration and glycolytic flux are major determinants of net acetate flux responds^7,10^. Both factors were therefore varied in *S. cerevisiae*. Increasing extracellular acetate from 5 to 60 mM did not increase acetate assimilation, and the net acetate flux remained close to zero (−0.1 ± 0.2 mmol·g_DCW_^−1^·h^−1^, Fig. 2C); higher acetate availability alone was thus insufficient to drive net assimilation.

**Fig. 2.**
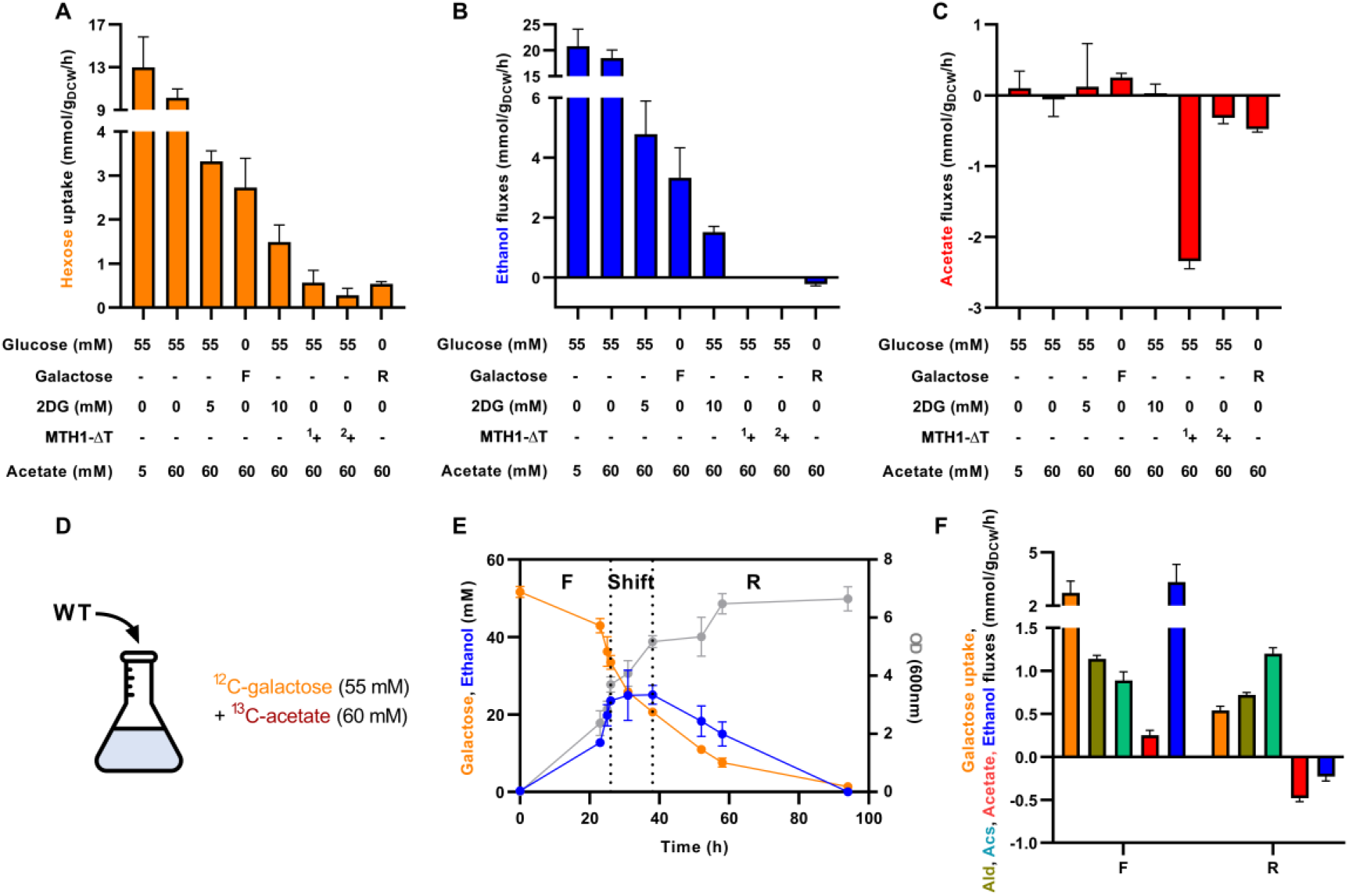
Net acetate flux is uncoupled from hexose uptake and shifts toward assimilation during respiratory metabolism. **(A)** Hexose uptake **(B)** net ethanol and **(C)** net acetate fluxes of *S. cerevisiae* grown in synthetic medium with 55 mM glucose or galactose and 5 or 60 mM [U-^13^C]acetate, supplemented where indicated with 5 or 10 mM 2DG. Galactose cultures were analyzed in the fermentative (F) and respiratory (R) phases, and *MTH1-ΔT* cultures in the first (^1^+) and second (^2^+) growth phases. **(D)** Experimental design for **(E, F)**: wild-type cells grown on 55 mM galactose with 60 mM [U-^13^C]acetate. **(E)** Galactose and ethanol concentrations and OD_600nm_ over time. Dotted lines delimit the fermentative (F), transition (Shift) and respiratory (R) phases. **(F)** Galactose uptake and net acetate and net ethanol fluxes, with Ald and Acs fluxes estimated by isotopic flux analysis, in the fermentative (F) and respiratory (R) phases. Fluxes are expressed in mmol·gDCW^−1^·h^−1^. Data are mean ± SD of 3 biological replicates; **p** values were calculated by Student’s *t*-tests. 2DG, 2-deoxyglucose.

Hexose uptake was next decreased using three complementary approaches: inhibition of glucose phosphorylation with 2-deoxyglucose (2DG)^22^, replacement of glucose with the more slowly consumed hexose galactose, and expression of *MTH1-ΔT*, which reduces glucose transporter expression and thereby glucose uptake even in the presence of high glucose^23^. Growth on galactose (Fig. 2D) comprised an initial fermentative phase with ethanol production, followed by a respiratory phase with ethanol consumption (Fig. 2E), allowing acetate exchange to be compared between the two metabolic states.

All three approaches decreased hexose uptake, although to different extents (Fig. 2A). Ethanol production closely tracked hexose uptake (r^2^ = 0.98, Fig. 2B and Supplementary Fig. 2), consistent with the known coupling between glycolytic flux and ethanol overflow^24,25^. By contrast, net acetate flux was essentially unrelated to hexose uptake (r^2^ = 0.02, Fig. 2C and Supplementary Fig. 3). Neither lowering glucose uptake with 2DG nor the fermentative phase on galactose resulted in net acetate assimilation. Net assimilation occurred only during the respiratory phase on galactose and, in its first growth phase, more strongly in the *MTH1-ΔT* strain growing on glucose (Fig. 2C, Supplementary Fig. 4), two conditions associated with reduced fermentation and increased respiratory metabolism^23,26^ (Fig. 2B).

**Fig. 3.**
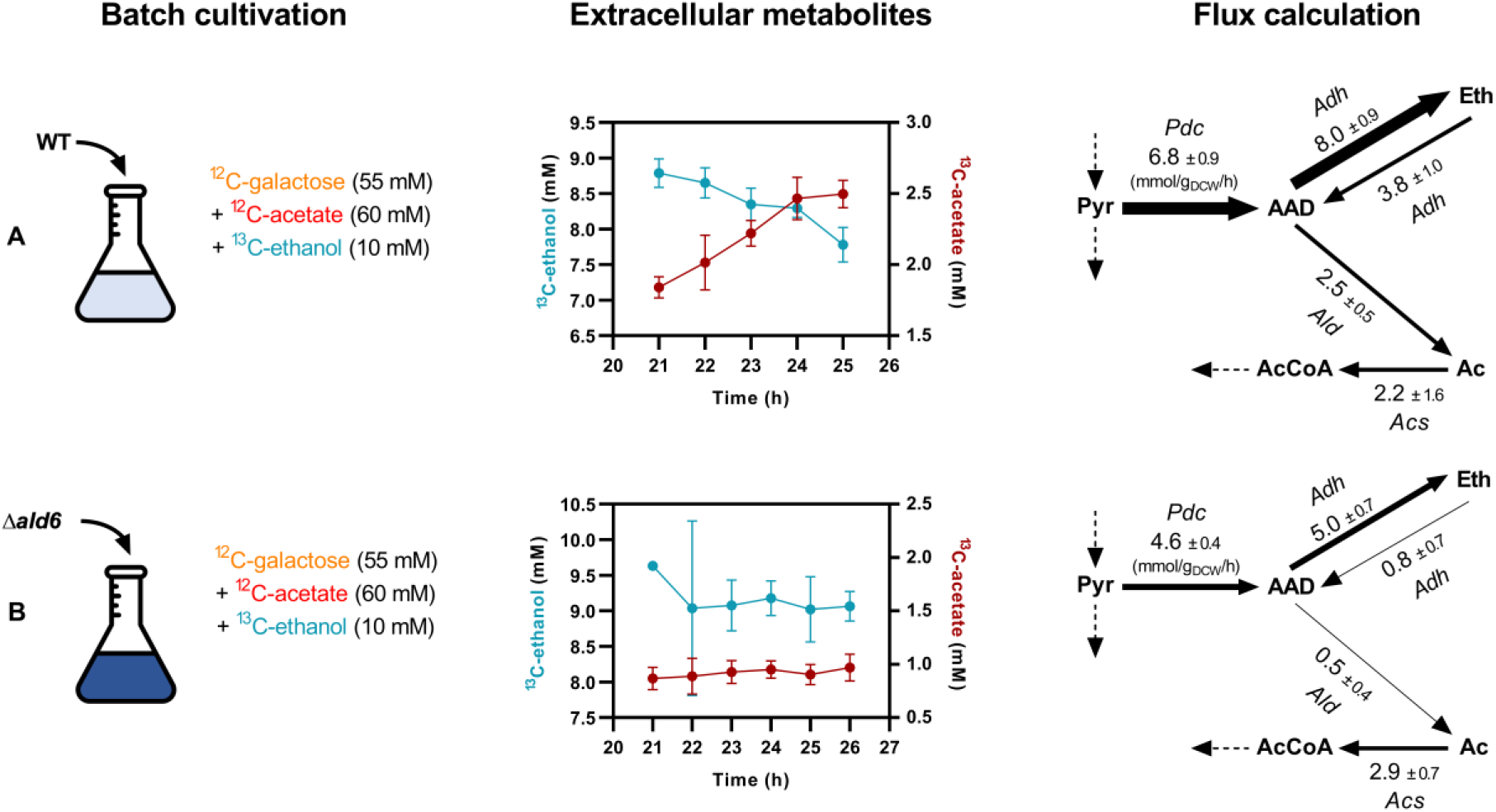
Extracellular ethanol is oxidized to acetate via Ald6 during fermentative growth. **(A)** Wild-type and **(B)** *Δald6* cells grown on 55 mM galactose with 60 mM unlabeled acetate and 10 mM [U-^13^C]ethanol. Left, experimental design. Middle, extracellular concentrations of ^13^C-ethanol and ^13^C-acetate during the fermentative phase. Right, fluxes (mmol·g_DCW_^−1^·h^−1^) through Pdc, Adh (forward and reverse), Ald and Acs estimated by isotopic flux analysis; arrow width is proportional to flux, and dashed arrows indicate connections to the rest of the network, whose fluxes are not shown. Data are mean ± [SD] of 3 biological replicates. Pyr, pyruvate; AAD, acetaldehyde; Eth, ethanol; Ac, acetate; AcCoA, acetyl-CoA; Pdc, pyruvate decarboxylase; Adh, alcohol dehydrogenase; Ald, aldehyde dehydrogenase; Acs, acetyl-CoA synthetase.

**Fig. 4.**
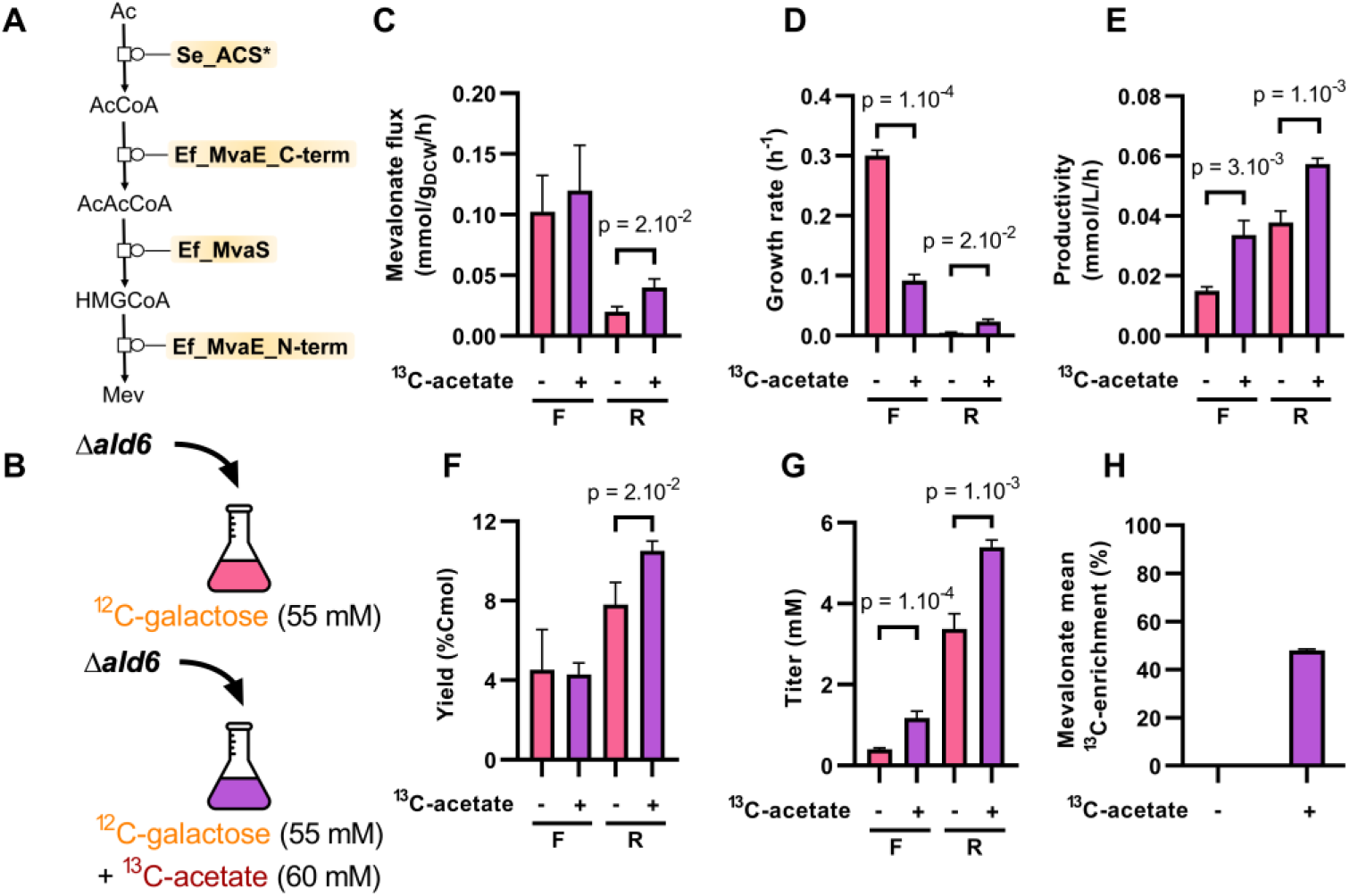
Extracellular acetate serves as a co-substrate for mevalonate production. **(A)** Acetate -to-mevalonate pathway expressed in the *S. cerevisiae Δald6* strain: Se_ACS* (acetyl-CoA synthetase), Ef_MvaE_C-term (acetoacetyl-CoA thiolase), Ef_MvaS (HMG-CoA synthase) and Ef_MvaE_N-term (HMG-CoA reductase). **(B)** Experimental design: the pathway expressing *Δald6* strain grown on 55 mM galactose with or without 60 mM [U-^13^C]acetate. **(C)** Mevalonate flux, **(D)** growth rate, **(E)** productivity, **(F)** yield and **(G)** titer in the fermentative (F) and respiratory (R) phases, without (−) or with (+) [U-^13^C]acetate. **(H)** Mean ^13^C-enrichment of mevalonate in the respiratory phase. Data are mean ± [SD] of 3 biological replicates; *P*-values were calculated by Student’s *t*-tests . Ac, acetate; AcCoA, acetyl-CoA; AcAcCoA, acetoacetyl-CoA; HMGCoA, hydroxymethylglutaryl-CoA; Mev, mevalonate.

Isotopic tracing and flux analysis revealed the processes underlying these different net responses. In 2DG cultures, assimilation of ^13^C-acetate and production of unlabeled acetate both persisted despite reduced glucose uptake, leaving the total acetate concentration essentially unchanged (Supplementary Fig. 5). In contrast, the transition from fermentative to respiratory growth on galactose shifted both ethanol and acetate from net production to net consumption (*p* = 4 × 10^−3^ and 6 × 10^−5^, respectively). For acetate, this shift reflected both a decrease in Ald flux and an increase in Acs flux (*p* = 1 × 10^−4^ and 1 × 10^−3^, respectively; Fig. 2F), not a change in either alone. Lowering glycolytic flux therefore does not by itself determine the direction of acetate exchange; net acetate assimilation instead results from concurrent decrease in production and increase in assimilation during the transition toward respiratory metabolism.

### Ethanol is oxidized to acetate during net ethanol production

The association between net acetate assimilation and loss of net ethanol production suggests that ethanol turnover contributes to acetate excretion. To test this, [U-^13^C]ethanol (10 mM) was traced during fermentative growth on 55 mM galactose with 60 mM extracellular acetate. Consistent with the known bidirectional turnover of ethanol^11^, cells consumed ^13^C-ethanol while simultaneously producing unlabeled ethanol from galactose (Supplementary Fig. 6). ^13^C-acetate was concurrently excreted (Fig. 3A), demonstrating that extracellular ethanol is oxidized to acetate even as cells remain net ethanol producers.

To assess the contribution of this route to net acetate exchange, we deleted *ALD6*, which encodes the main cytosolic aldehyde dehydrogenase responsible for acetate formation^17^. In the *Δald6* strain, ^13^C-ethanol remained essentially constant and excretion of ^13^C-acetate was strongly reduced, whereas unlabeled ethanol accumulated in the medium (Fig. 3B). The reverse Adh and the Ald fluxes were respectively 4.7- and 5.3-fold lower than in the wild-type strain, but acetate assimilation through Acs was maintained (Fig. 3A,B). As a result, net acetate flux shifted to assimilation (−2.4 ± 0.5 mmol·g_DCW_^−1^·h^−1^; Supplementary Fig. 7). This shift did not require exogenous ethanol: without added ethanol the *Δald6* strain still assimilated extracellular acetate during the fermentative phase (−1.8 ± 0.5 mmol·g_DCW_^−^1·h^−1^), with Ald flux decreasing from 1.14 ± 0.04 to 0.41 ± 0.02 mmol·g_DCW_^−1^·h^−1^, and an increase in Acs flux, from 0.89 ± 0.10 to 2.18 ± 0.44 mmol·g_CCW_^−1^·h^−1^ (Supplementary Fig. 8). Ald6-dependent acetate production, to which extracellular ethanol oxidation contributes, therefore determines the direction of net acetate exchange during fermentative growth.

### Increased acetyl-CoA demand channels extracellular acetate into bioproduction

The substantial acetate assimilation observed here suggests that extracellular acetate can constitute an alternative source of carbon for biosynthesis during fermentative and respiratory growth. A synthetic acetate-to-mevalonate pathway was therefore expressed to test whether increasing the metabolic demand for acetate-derived acetyl-CoA redirects acetate toward assimilation. The pathway involved three enzymes, encoded by four polypeptides: Acs^L641P^ from *Salmonella enterica* (Se_ACS*), MvaS from *Enterococcus faecalis* (Ef_MvaS), and MvaE from *E. faecalis*, the latter expressed as two separate polypeptides (Ef_MvaE_C-term and Ef_MvaE_N-term) (Fig. 4A).

This pathway was expressed in the *Δald6* strain, which has reduced endogenous acetate production, and the cells were grown on galactose with [U-^13^C]acetate (Fig. 4B). This combination provides an additional biosynthetic sink for acetate-derived acetyl-CoA, allowing a direct test of whether extracellular acetate can be redirected into product.

During the fermentative phase, expression of the acetate-to-mevalonate pathway did not measurably increase acetate assimilation (Supplementary Fig. 9). Nevertheless, acetate supplementation increased mevalonate titer (+195%) and productivity (+127%) despite slower growth (−69%; Fig. 4G,E,D), indicating that the benefits of redirecting carbon toward mevalonate synthesis outweigh the fermentative-phase growth cost. During the respiratory phase, expression of the pathway increased Acs flux and acetate assimilation (Supplementary Fig. 10). Under these conditions, acetate supplementation increased the mevalonate flux (+100%, Fig. 4C), growth rate (+389%, Fig. 4D), productivity (+50%, Fig. 4E), yield (+34%, Fig. 4F) and titer (+59%, Fig. 4G). Isotopic analysis confirmed incorporation of extracellular ^13^C-acetate into mevalonate, with 48% of mevalonate carbon derived from acetate in the respiratory phase (Fig. 4H). Acetate therefore did not merely alter mevalonate production, but directly provided a substantial fraction of the carbon incorporated into the product.

Increasing the demand for mevalonate biosynthesis can thus enhance acetate assimilation and channel extracellular acetate into an acetyl-CoA-derived product. Rather than solely accumulating as an overflow metabolite, extracellular acetate can therefore serve as a productive co-substrate for bioproduction.

## Discussion and conclusion

Extending recent work that has challenged the view of overflow metabolites as metabolic waste^7,10–13^, this study shows that acetate in *S. cerevisiae* can be recycled as a nutrient and influence cell physiology. During fermentative and respiratory growth, acetate excretion and assimilation proceed simultaneously at fluxes more than an order of magnitude larger than the net difference. Extracellular acetate overflow therefore reflects only the imbalance between these fluxes, not their magnitude: even under conditions classically interpreted as acetate production during glucose consumption followed by reassimilation after glucose depletion, acetate is in fact being excreted and assimilated at the same time. Acetate overflow is thus not a one-way process but the net result of opposing production and assimilation fluxes.

In fermenting yeast, acetate exchange is metabolically connected to ethanol turnover. Xiao et al. showed that extracellular ethanol is rapidly reassimilated and provides a substantial source of cytosolic and mitochondrial acetyl-CoA^11^. The present results resolve the link between acetate and ethanol metabolism: during hexose fermentation, ethanol oxidation contributes directly to extracellular acetate excretion, while acetate is simultaneously assimilated into acetyl-CoA metabolism. Ethanol and acetate overflow are therefore not independent outputs of hexose metabolism; rather, acetate exchange links ethanol oxidation to acetyl-CoA utilization, allowing ethanol-derived carbon released from the cell to return to central metabolism via extracellular acetate.

Pathway organization shapes acetate exchange differently in *E. coli* and *S. cerevisiae*. In *E. coli*, acetate excretion and assimilation during glucose metabolism are mostly mediated by the same reversible Pta-AckA pathway, whose net flux depends strongly on the extracellular acetate concentration and the glycolytic flux^7,10^. In yeast, by contrast, acetate is produced by aldehyde dehydrogenase and assimilated by acetyl-CoA synthetase, so production and assimilation proceed through separate, irreversible routes. Consistent with this structure, a higher extracellular acetate concentration did not substantially increase assimilation. In *E. coli*, extracellular acetate concentration and glycolytic flux largely dictate acetate flux; in yeast, neither does because production and assimilation are catalyzed by separate, independently regulated enzymes.

These results further decouple carbon entry flux from acetate flux. In *S. cerevisiae*, ethanol production closely follows hexose uptake^24,27^, whereas here, net acetate flux did not. The direction of acetate exchange shifted toward assimilation as metabolism became respiratory, acetate production decreasing through lower Ald flux and acetate assimilation increasing through higher Acs flux. Deletion of *ALD6* shifted the balance toward assimilation in the same way. This dependence on the metabolic state downstream of glycolysis may also explain the few previous reports of acetate co-utilization in yeast, notably during aerobic xylose metabolism, an engineered condition characterized by weak ethanol overflow and active respiration^19^.

Rather than establishing acetate co-utilization as a general feature of yeast metabolism, these observations identify specific conditions in which acetate exchange shifts toward recovery: cells with similarly low glycolytic flux can show opposite net acetate fluxes depending on their metabolic state. Net acetate flux is set by the balance between distinct production and assimilation fluxes, which respond differently to metabolic perturbations. The present experiments do not isolate a single causal signal – respiration itself, relief of glucose repression or another regulatory mechanism – but they show that the direction of acetate exchange emerges downstream of glycolysis, from coordinated changes in ethanol turnover, acetate production and acetyl-CoA utilization.

This coupling of acetate exchange to acetyl-CoA metabolism may itself be physiologically significant. Acetate assimilation increased both during respiratory metabolism and upon addition of a biosynthetic sink for acetate-derived acetyl-CoA. Extracellular acetate therefore constitutes a recoverable source of acetyl-CoA carbon even as acetate is produced. Acetate exchange may help buffer mismatches between acetyl-CoA supply and utilization by allowing carbon to be transiently released and subsequently recovered. These results do not demonstrate that acetate contributes directly to acetyl-CoA homeostasis but they do reveal the flux organization that would enable it.

In summary, this study provides a quantitative framework for when acetate becomes a co-substrate in yeast. What appears, from extracellular concentrations alone, to be acetate overflow is the net result of simultaneous production and assimilation, whose balance shifts toward assimilation with a respiratory metabolic state, reduced acetate production, or increased demand for acetate-derived acetyl-CoA. Exploiting the last of these, extracellular acetate carbon was routed into mevalonate production. Acetate co-utilization therefore emerges not by acetate assimilation *de novo*, but by shifting a pre-existing exchange from net release toward recovery.

## Materials and methods

### Plasmid construction

The plasmids used in this study (Supplementary Table 1) were constructed by Golden Gate assembly using the MoClo Kit^27,28^ and yeast homologous recombination. The primers (Supplementary Table 2) were synthesized by IDT (Leuven, Belgium) and the coding sequences of Se_Acs^L641P^, Ef_MvaS, Ef_MvaE_N-term, Ef_MvaE_C-term and MTH1-ΔT (Supplementary Table 3), codon-optimized for *S. cerevisiae*, by Twist Bioscience (San Francisco, California).

The acetate mevalonate pathway used in this study was constructed using three genes: Se_Acs^L641P^ from *Salmonella enterica*, Ef_MvaS from *Enterococcus faecalis* and Ef_MvaE from *E. faecalis*, the latter expressed as separate N- and C-terminal domains expressed as separate polypeptides (Ef_MvaE_N-term and Ef_MvaE_C-term). Four transcriptional units were constructed by Golden Gate assembly using BsaI-HF and a T4 DNA Ligase (New England Biolabs) generating the vectors: pI2M013, pI2M017, pI2M018 and pI2M021. Promoters, ORFs and terminators were assembled over 25 cycles of 1 min at 37°C and 1 min at 16°C, followed by 5 min at 60°C.

The final vector, pI2M114, carrying the four transcriptional units, was assembled by yeast homologous recombination using the Golden Gate assembled vectors: pI2M013, pI2M017, pI2M018, and pI2M021, together with pMYT059 and linearized pRS316^29^. pRS316 was linearized by PCR with primers oI2M139 and oI2M140 using KOD Xtreme polymerase (Sigma-Aldrich) and gel-purified with the Monarch DNA Gel Extraction Kit (New England Biolabs); the transcriptional-unit vectors were linearized with NotI (Thermo Fisher Scientific, Lithuania). After co-transformation into yeast, the assembled plasmid was verified by colony PCR.

For plasmid rescue, positive clones were grown overnight in 10 mL SD-URA. The culture was centrifuged (10 min, 4000 × g) and the pellet was resuspended in 1 mL of TE buffer (100 mM Tris-HCl 1 mM EDTA pH 8). After a further centrifugation, the cell pellet was resuspended in 200 µL of Y1 buffer (1 M Sorbitol, 100 mM EDTA, 10 mM DTT), the cell wall was digested with 10 µL of lyticase (8.5 U/µL in 40% glycerol) for 1 h at 37°C and the cell suspension was centrifuged (10 min, 4000 × g). Plasmid was extracted from the pellet with the EZ-10 Spin Column Plasmid DNA Miniprep Kit (BioBasic), eluted in 20 µL of TE buffer at 55°C for 10 min, transformed into NEB 5-alpha competent *E. coli* (New England Biolabs) and sequenced by Sanger sequencing (Eurofins).

The integrative vector pI2M168 was constructed by assembling pMYT075 (homology arms for locus I) with pMYT031 (*HIS3* marker) by BbsI Golden Gate assembly. The weak *RAD27* promoter, *MTH1-ΔT* — carrying an internal 225-bp deletion (nucleotides 169–393 of the *MTH1* ORF) — and the *ADH1* terminator were then cloned into this backbone by BsaI Golden Gate assembly, yielding pI2M169, which was integrated at locus I to overexpress *MTH1-ΔT*.

### Strain construction

All strains used in this study are derivatives of *S. cerevisiae* CEN.PK2-1D (MATα, ura3-52, trp1-289, leu2-3,112, his3Δ1) and are listed in Supplementary Table 4. Yeast cells were transformed using the high-efficiency lithium acetate protocol^30^. The *ALD6* gene was deleted using the short flanking homology method^31^: the *natMX* cassette was amplified from pUG74 with the primers listed in Supplementary Table 2 and integrated by homologous recombination. This deletion was confirmed by colony PCR.

### Growth conditions and media composition

All cultures were grown at 28°C with shaking at 200 rpm in an INFORS HT TR-225 incubator.

*S. cerevisiae* was grown in YPD (yeast extract/peptone/glucose) prior to transformation; after transformation, strains were selected on auxotrophic markers or 250 µg/mL nourseothricin (Jena Bioscience).

For characterization, cells were grown in synthetic medium (pH 5.5) containing yeast nitrogen base (YNB, 6.9 g/L; Formedium) and CSM-URA (0.67 g/L; MP Biomedicals), supplemented with combinations of carbon sources: glucose (Formedium), galactose (Formedium), sodium acetate (AnalaR NORMAPUR, VWR/Prolabo), U-^13^C-acetate (Eurisotop, France), and U-^13^C-ethanol (Cambridge Isotope laboratories); concentrations are given in the Results section. Fresh colonies were inoculated into 48-well culture plates containing 350 µL of SC-dropout medium with 2% (w/v) glucose and grown overnight. These precultures were used to inoculate 20 mL of synthetic medium in 100 mL shake flasks to an initial OD_600_ of 0.05, except the 10 mM 2-deoxyglucose (2DG), which was started at an OD_600_ of 0.2.

Optical density was measured at 600 nm with a PowerWave spectrophotometer (BioTek Instruments). Biomass concentration was calculated from OD_600_ as biomass (g_DCW·_L^−1^) = OD_600_ / 3.416^32^.

## Analytical methods

Glucose, galactose, ethanol, glycerol, mevalonate, and acetate were quantified by high-performance liquid chromatography (HPLC). Culture supernatants were collected by centrifugation (10 min, 13,000 × g) and transferred into vials. Separation was performed on an Ultimate 3000 system (Thermo Fisher Scientific, Lithuania) equipped with a Bio-Rad Aminex HPX-87H column (300 × 7.8 mm, 9 µm) and the RI-101 refractive-index detector (Shodex), at 60°C with 5 mM H_2_SO_4_ as the mobile phase at 0.6 mL·min^−1^. Chromatograms were analyzed with Chromeleon software (Thermo Fisher Scientific).

For the labeling experiments, [U-^13^C]acetate (sodium acetate-U-^13^C_2_, 99%) or [U-^13^C]ethanol (ethanol-U-^13^C_2_, 99%, <6% H_2_O) was added to the medium. Supernatant (180 µL), collected as described above, was mixed with 20 µL of internal standard (2.35 g·L^−1^ TSP-d4 in D_2_O). ^1^H-NMR spectra were acquired on an Avance III 500 MHz spectrometer with a 5-mm z-gradient BBI probe (Bruker, Germany).

## Mass spectrometry

The ^13^C-enrichment of mevalonate was measured by ion exchange liquid chromatography, using a Thermo Scientific Dionex ICS-5000+ Reagent-Free HPIC system coupled to a Thermo Scientific Q Exactive Plus hybrid quadrupole-Orbitrap mass spectrometer equipped with a heated electrospray ionization probe (HESI-II)^33^. Mevalonate isotopologue intensities were extracted from the raw data using Skyline v24.1.0 and the mean ^13^C-enrichment was calculated from the isotopologue distribution with IsoCor v2.2.2^34^ (https://github.com/MetaSys-LISBP/IsoCor), after correcting for naturally occurring isotopes and the isotopic purity (99%) of the ^13^C-labeled acetate.

## Data analysis

The model developed to quantify metabolic fluxes consists of 20 reactions, 20 species, and 2 compartments (the environment and the cell; Fig. 1A).

The differential equations, which balance the concentrations of extracellular compounds (biomass, glucose, glycerol, ethanol, and acetate) and intracellular compounds (glyceraldehyde-3-phosphate, pyruvate, acetaldehyde, and acetyl-CoA), were augmented with isotopic equations for parameter estimation. As described previously^35^, all reactions except biomass synthesis were considered separately for unlabeled and labeled reactants. Fluxes were assumed to be constant over time (metabolic steady-state assumption).

The final model has 14 free parameters in total. Model parameters (p) were estimated by fitting the measured concentrations of biomass, glucose, glycerol, and unlabeled and labeled ethanol and acetate, minimizing the objective function *f*, the weighted sum of squared errors:

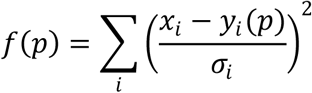

where *x*_*i*_ is the experimental value of data point *i*, with an experimental standard deviation *σ*_*i*_, and *y*_*i*_(*p*) is the corresponding simulated value. *f* was minimized by particle swarm optimization (4,000 iterations, swarm size of 50).

The model was built and analyzed using COPASI^36^ (v4.39) and is provided in SBML and COPASI formats in the Supplementary Material and at https://github.com/MetaSys-LISBP/Yeast_acetate along with the data for all replicates.

Mevalonate fluxes were calculated from mevalonate and biomass concentration-time profiles with PhysioFit^37^ (https://github.com/MetaSys-LISBP/PhysioFit).

## Statistical analysis

Comparisons between strains or conditions were performed using Student’s *t*-tests in GraphPad Prism 8.3.

## Acknowledgments

We thank all members of the Molecular and Metabolic Team at the Toulouse Biotechnology Institute for their support and for fostering an excellent research environment. The authors thank MetaboHub-MetaToul (Metabolomics & Fluxomics facilities, Toulouse, France, https://mth-metatoul.com), which is part of the French National Infrastructure for Metabolomics and Fluxomics (https://www.metabohub.fr), funded by the ANR (MetaboHUB-ANR-11-INBS-0010), for access to MS facilities. SO Ferm platform, more particularly Alicia Huesca, for access and help with the HPLC. We are also grateful to Dr. Ellis and Dr. Dueber for providing the plasmids used in this study. This work was supported by Carnot Institute 3BCAR (AAP 2024 AmiSynpa), by a government grant managed by the Agence Nationale de la Recherche under the Investissements d’Avenir programme (ANR-18-EURE-0021), and by the MITI program of CNRS (METABO-FLOW grant).

## Author contributions

Funding acquisition: PM, SCC; Project administration: SCC; Conceptualization: AC, PM, SCC; Investigation: AC, MZK, SCC; Formal analysis: AC, PM, SCC; Software: AC, PM; Resources: HK; Supervision: HMY, SCC; Visualization: AC, PM, SCC; Writing– original draft: AC, PM, SCC; Writing – review & editing: AC, MZK, HK, GT, HMY, PM, SCC.

