## Supplementary figures and tables for "Bidirectional acetate exchange couples ethanol turnover to acetyl-CoA metabolism in *Saccharomyces cerevisiae*"

Carneiro et al.

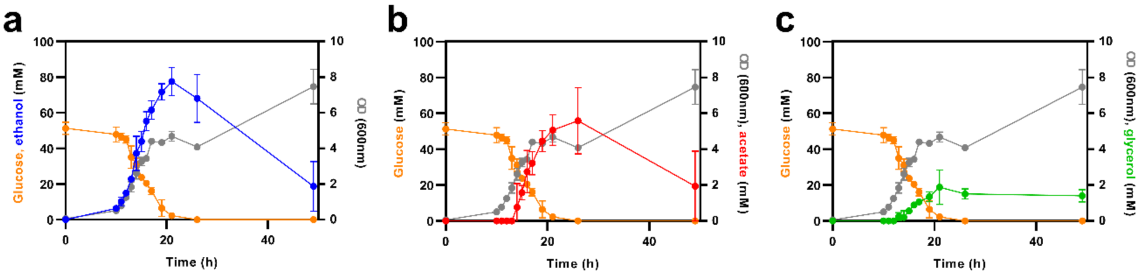


**<u>Supplementary Figure 1.</u>** Growth of *S. cerevisiae CEN.PK2-1D* in synthetic medium with 55 mM glucose. Time course of **(a)** ethanol **(b)** acetate and **(c)** glycerol concentrations. Optical density at 600 nm is represented in grey and glucose concentration in orange.

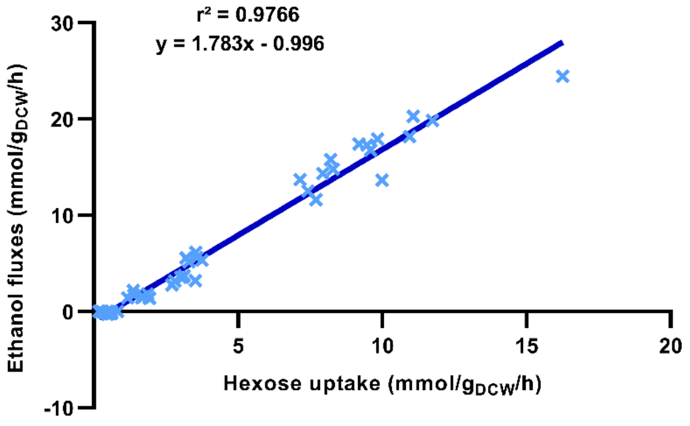


**Supplementary Figure 2.** Correlation between hexose uptake and ethanol fluxes. Data were pooled from all experiments performed in this study. Each cross represents one biological replicate and one culture condition.

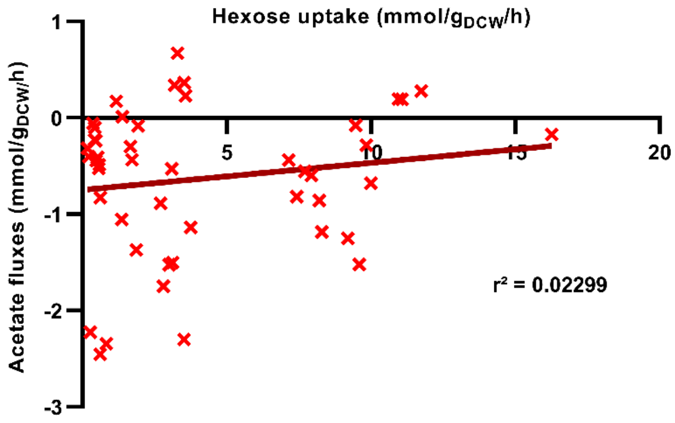


**<u>Supplementary Figure 3.</u>** Correlation between hexose uptake and acetate fluxes. Data were pooled from all experiments performed in this study. Each cross represents one biological replicate and one culture condition.

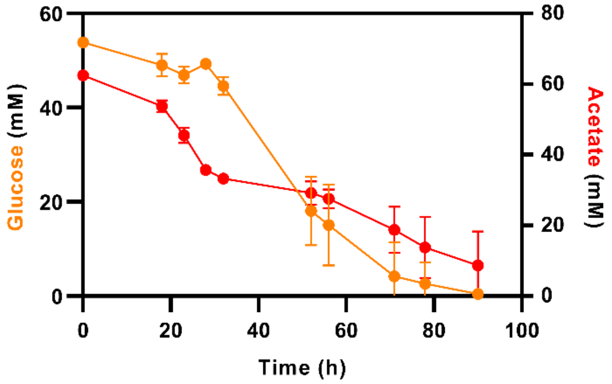


**<u>Supplementary Figure 4.</u>** Growth of the *S. cerevisiae MTH1-ΔT* strain in synthetic medium with 55 mM glucose and 60 mM acetate. Time course of glucose (orange) and acetate (red) concentrations.

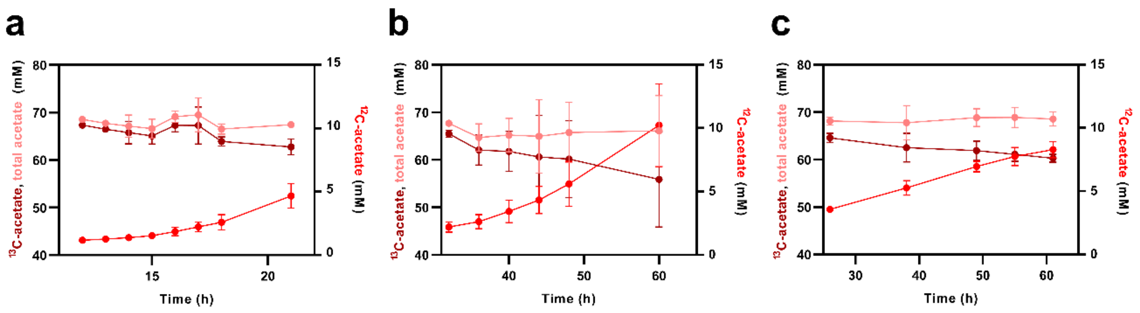


**<u>Supplementary Figure 5.</u>** Time course of differently labeled acetate species during growth on synthetic medium with 55 mM glucose and 60 mM acetate. $^{12}C$-acetate (bright red), $^{13}C$-acetate (dark red) and total acetate (light pink) were measured in the wild-type strain with 0 **(a)**, 5 **(b)** or 10 mM of 2DG **(c).**

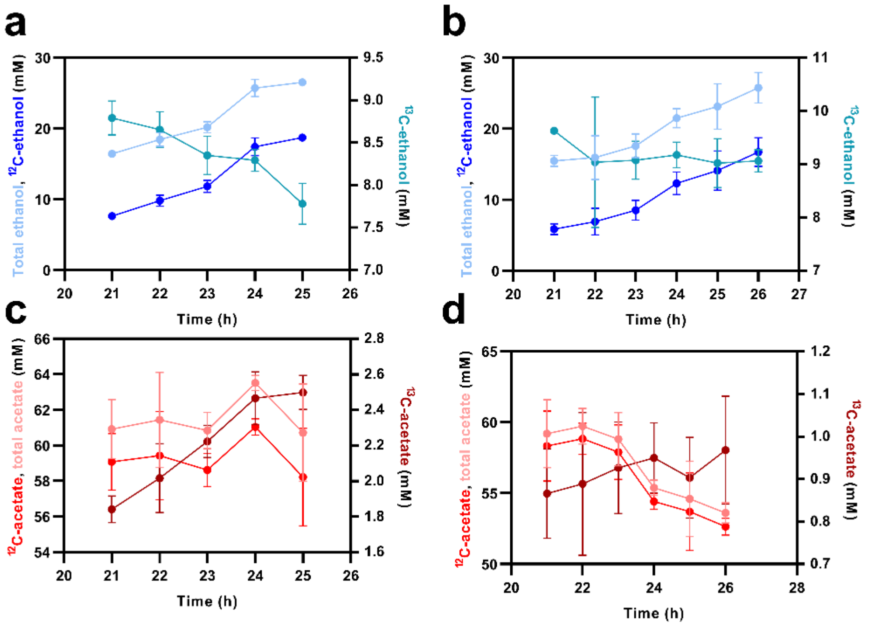


**<u>Supplementary Figure 6.</u>** Time course of differently labelled ethanol and acetate species during growth on synthetic medium with 55 mM galactose, 60 mM $^{12}$C-acetate and 10 mM $^{13}$C-ethanol in fermentative phase. $^{12}$C-ethanol (bright blue), $^{13}$C-ethanol (turquoise blue) and total ethanol (pale blue) were measured in the wild-type strain **(a)** and the Δ*ald6* strain **(b)**. $^{12}$C-acetate (bright red), $^{13}$C-acetate (dark red) and total acetate (pale red) were measured in the wild-type strain **(c)** and the Δ*ald6* strain ***(d)***.

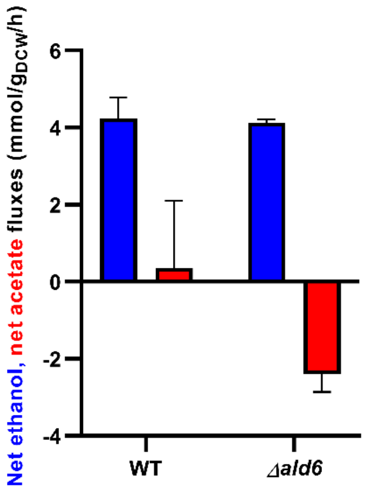


**Supplementary Figure 7.** Effect of *ALD6* deletion on net ethanol and acetate fluxes. Net ethanol and acetate fluxes in wild-type and Δ*ald6* strains grown in synthetic medium with 55 mM galactose, 60 mM $^{12}$C-acetate and 10 mM $^{13}$C-ethanol in fermentative phase.

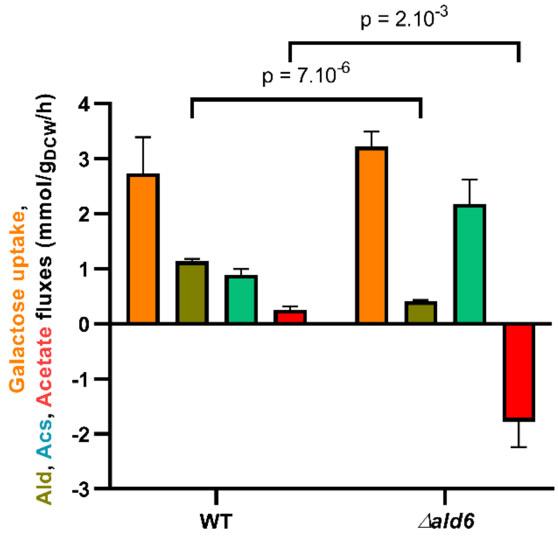


**<u>Supplementary Figure 8.</u>** Effect of *ALD6* deletion on acetate overflow related fluxes. Galactose uptake and Ald, Acs and net acetate fluxes in *S. cerevisiae* wild-type and Δ*ald6* strains grown in synthetic medium containing with 55 mM galactose and 60 mM acetate in fermentative phase.

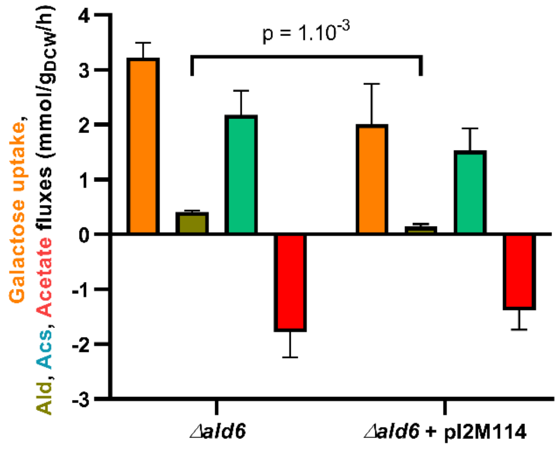


**Supplementary Figure 9.** Effect of synthetic mevalonate pathway expression on acetate overflow related fluxes during the fermentative phase. Galactose uptake rate and Ald, Acs and net acetate fluxes in *S. cerevisiae* Δ*ald6* strains with or without pI2M114 grown in synthetic medium containing 55 mM galactose and 60 mM of $^{13}$C-acetate in the fermentative phase.

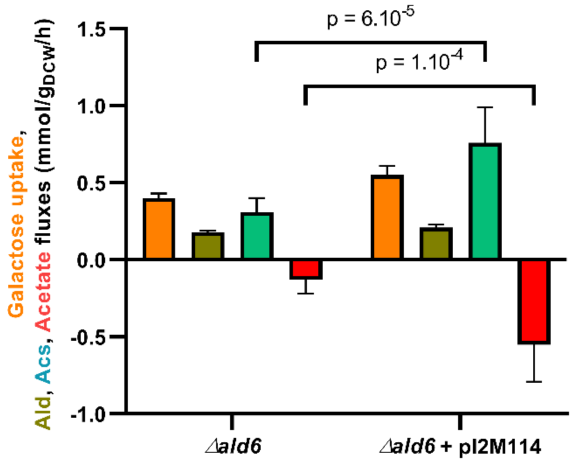


**<u>Supplementary Figure 10.</u>** Effect of synthetic mevalonate pathway expression on acetate overflow related fluxes during the respiratory phase. Galactose uptake rate and Ald, Acs and net acetate fluxes in *S. cerevisiae* Δ*ald6* strains with or without pI2M114 grown in synthetic medium containing 55 mM galactose and 60 mM of $^{13}$C-acetate in the respiratory phase.

**<u>Supplementary Table 1 :</u>** The list of plasmids used in this study

| Name | Description | Source |
|---|---|---|
| pRS316 | CEN6/ARS4-URA3; AmpR; ColE1 | [1] |
| pUG74 | loxP-natMX6-loxP; AmpR; ColE1 | [2] |
| pMYT031 | His3, CamR | [3] |
| pMYT039 | S/1 Assembly Cassette, AmpR | [3] |
| pMYT043 | 2/3 Assembly Cassette, AmpR | [3] |
| pMYT045 | 3/4 Assembly Cassette, AmpR | [3] |
| pMYT046 | 4/E Assembly Cassette, AmpR | [3] |
| pMYT059 | 1/2 Spacer Cassette, AmpR | [3] |
| pMYT075 | Int. 1 Vector, KanR | [3] |
| pYTK009 | pTDH3, CamR | [4] |
| pYTK010 | pCCW12, CamR | [4] |
| pYTK011 | pPGK1, CamR | [4] |
| pYTK012 | pHHF2, CamR | [4] |
| pYTK025 | pRAD27, CamR | [4] |
| pYTK053 | tADH1, CamR | [4] |
| pYTK054 | tPGK1, CamR | [4] |
| pYTK055 | tENO2, CamR | [4] |
| pYTK056 | tTDH1, CamR | [4] |
| pI2M013 | pMYT039, pYTK010, AcsL641P, pYTK055 | This study |
| pI2M017 | pMYT043, pYTK011, MvaE_N-term, pYTK056 | This study |
| pI2M018 | pMYT045, pYTK009, MvaS, pYTK053 | This study |
| pI2M021 | pMYT046, pYTK012, MvaE_C-term, pYTK054 | This study |
| pI2M114 | pRS316 linearized, URA3 marker, Se-acsL641P, Ef-MvaS, Ef-MvaE_N-term, Ef-MvaE_C-term | This study |
| pI2M168 | pMYT075, pMYT031 | This study |
| pI2M169 | pI2M168, His3 marker, MTH1-ΔT | This study |

**Supplementary Table 2.** The list of oligonucleotides used in this study.

| Name | Primer sequence | Position/ATG |
|---|---|---|
| *ALD6* in pUG74 | ATGACTAAGCTACACTTTGACACTGCTGAACCAGTCAAGATC**agctgaagcttcgtacgc** | 0 to +42 |
| | TTACAACTTAATTCTGACAGCTTTTACTTCAGTGTATGCA**ataggccactagtggatctg** | +1463 to +1503 |
| oI2M139 | gaaatctgctcgtcagtggtgctcacactgacgaatcatgtacagatcatAGCTTATCGATACCGTCGAC | / |
| oI2M140 | GTATTGCGACGAATTGCCACGTTGTCGCAGTGTCTGTACTGTGGGTTAATGGGGATCCACTAGTTCTAGAGC | / |

Sequence complementary to pUG74 is indicated in bold type

**Supplementary Table 3.** The list of gene sequences used in this study.

| Name | Gene sequence |
|---|---|
| AcsL641P | GgctacGGTCTCAtatgATGAGCCAGACGCACAAGCACGCCATACCTGCTAACATCGCTGACCGTTGTTTGATCAATCCGGAGCAATATGAGACAAAATACAAACAATCGA<br>TAAATGACCCCGACACATTTTGGGGCGAACAAGGAAAGATTCTAGATTGGATAACTCCCTACCAGAAAGTAAAGAATACGTCTTTCGCCCCCGGTAACGTGTCCATAAA<br>GTGGTACGAGGACGGTACGCTGAATTTAGCAGCCAATTGTTTAGACAGGCATCTTCAGGAGAACGGAGACAGGACAGCTATAATATGGGAAGGAGATGATACTTCACA<br>GTCGAAGCACATTTCCTATCGTGAATTACATAGGGACGTGTGCCGTTTCGCCAACACCCTTTTAGACCTGGGCATTAAGAAAGGAGATGTTGTGGCCATCTATATGCCT<br>ATGGTCCCGGAAGCCGCCGTGGCTATGCTGGCGTGCGCAAGGATTGGCGCCGTACACAGTGTTATCTTCGGCGGTTTCAGTCCAGAGGCAGTTGCTGGAAGGATAAT<br>CGACTCGTCTAGCAGGTTAGTCATAACGGCCGACGAGGGAGTCAGGGCAGGTAGATCCATTCCATTGAAGAAGAATGTTGATGACGCGTTAAAGAATCCTAATGTAAC<br>TTCTGTAGAGCACGTTATAGTCTTAAAGAGAACCGGATCTGACATAGACTGGCAAGAAGGTAGGGATCTATGGTGGAGAGACTTGATAGAGAAAGCTAGTCCTGAGCA<br>CCAACCCGAGGCTATGAACGCTGAGGACCCATTGTTCATTCTATATACCAGTGGATCTACGGGGAAGCCTAAGGGAGTATTACACACTACCGGCGGTTACTTAGTGTA<br>TGCCGCTACTACTTTCAAGTACGTTTTCGACTATCATCCGGGAGACATTTACTGGTGCACGGCCGATGTGGGTTGGGTCACTGGCCACAGTTATCTATTATATGGTCCC<br>TTGGCGTGCGGTGCCACTACCTTAATGTTTGAGGGCGTGCCAAACTGGCCGACACCTGCCAGGATGTGTCAGGTGGTCGACAAGCACCAGGTCAATATTCTTTACAC<br>CGCTCCGACTGCGATTCGTGCTTTAATGGCAGAAGGGGACAAAGCTATAGAGGGTACTGACCGTAGTTCTCTAAGAATCTTGGGCAGTGTTGGTGAGCCAATAAATCC<br>GGAAGCGTGGGAGTGGTATTGGAAGAAGATTGGTAAGGAAAAGTGCCCGGTTGTGGACACATGGTGGCAGACAGAGACAGGTGGGTTCATGATAACACCACTACCC<br>GGAGCTATTGAACTGAAAGCCGGTTCTGCTACCAGACCTTTCTTTGGTGTCCAACCCGCACTGGTCGATAATGAGGGCCACCCCCAGGAAGGTGCAACCGAGGGAAA |
| MvaS | GgctacGGTCTCAtatgACCATCGGAATAGACAAGATATCGTTCTTCGTACCCCCGTATTATATTGACATGACAGCCCTAGCTGAGGCTAGAAATGTCGACCCAGGGAAGTT<br>TCATATAGGCATTGGCCAAGATCAGATGGCAGTCAATCCGATAAGTCAGGACATAGTTACATTCGCGGCCAACGCCGCCGAGGCAATACTTACGAAGGAAGATAAAGA<br>GGCTATTGACATGGTAATTGTGGGGACTGAGTCGTCCATTGACGAGTCAAAGGCTGCCGCAGTCGTATTGCATAGGCTAATGGGAATTCAACCATTTGCCAGGAGTTT<br>TGAGATTAAGGAAGCATGCTATGGGGCTACGGCAGGCTTACAATTGGCAAAGAATCATGTAGCATTACATCCAGATAAGAAGGTTCTGGTTGTAGCAGCAGACATAGC<br>CAAGTATGGACTGAACAGTGGTGGTGAGCCGACGCAGGGTGCAGGCGCGGTTGCCATGCTAGTTGCATCCGAACCTAGAATATTAGCTTTAAAAGAGGATAACGTCA<br>TGTTGACCCAGGATATTTATGACTTTTGGCGTCCAACAGGACATCCCTACCCAATGGTGGACGGGCCGTTATCTAACGAAACCTACATACAGTCATTCGCTCAGGTATG<br>GGATGAGCACAAGAAGAGAACGGGCTTGGATTTTGCGGATTATGACGCTTTAGCATTCCATATCCCTTACACTAAAATGGGAAAGAAGGCTCTATTAGCAAAGATCAGC<br>GATCAGACCGAGGCTGAACAGGAGAGGATCTTGGCCAGATACGAGGAATCTATTATTTACTCCAGACGTGTCGGAAACCTGTATACGGGGTCATTGTATTTGGGCTTG<br>ATTAGCCTTCTAGAAAATGCTACCACTTTAACTGCAGGCAATCAGATTGGCTTGTTCTCTTACGGTTCAGGCGCCGTAGCAGAGTTCTTCACGGGCGAGCTGGTTGCG<br>GGGTACCAGAACCACTTGCAAAAGGAAACGCACTTGGCCCTGCTTGACAACCGTACAGAATTATCTATCGCCGAATACGAGGCAATGTTTGCTGAAACATTAGATACTG<br>ACATCGACCAGACTCTTGAGGACGAGTTGAAGTATAGTATTAGTGCGATCAACAACACTGTAAGGTCCTACAGAAATGGTGGCGGTGGAAGCGGTGGTGGTGGAAGC<br>CTCGAGTACCCTTACGATGTTCCCGACTATGCCtaaATCCTGAGACCgtagcC |
| MvaE_C-term | GgctacGGTCTCAtatgAAGAACTCCAGGTTTTACCAGATGTCTCCGGAAGAAAGGCTTGCGAGTCTTTTGAACGAGGGTCAAATCAGCGCAGATACAAAGAAGGAATTTG<br>AGAATACCGCGCTTAGTTCACAAATTGCGAATCATATGATTGAGAATCAGATTAGTGAAACCGAGGTTCCTATGGGCGTGGGATTGCACCTGACGGTCGACGAGACAG<br>ACTACTTGGTGCCCATGGCAACAGAGGAACCATCTGTCATTGCTGCCCTGTCGAACGGTGCAAAGATAGCACAGGGTTTCAAAACGGTAAATCAGCAGCGTCTGATGA<br>GAGGACAGATTGTGTTCTACGACGTCGCCGATGCGGAATCACTGATAGACGAGCTACAGGTCAGGGAGACTGAGATTTTCCAACAGGCAGAGCTATCGTATCCTTCCA<br>TTGTAAAGAGAGGTGGTGGACTAAGGGACCTGCAGTATAGAGCTTTTGACGAGTCATTCGTCAGTGTAGACTTCTTAGTCGATGTGAAGGACGCTATGGGCGCGAACA<br>TCGTCAATGCCATGCTTGAGGGAGTTGCCGAGTTGTTTAGAGAGTGGTTCGCAGAGCAGAAAATCTTATTCAGCATTTTATCCAACTACGCAACCGAGTCTGTGGTAAC<br>AATGAAAACCGCCATACCTGTTAGCCGTCTTAGTAAGGGTTCAAATGGTAGGGAGATAGCAGAGAAAATAGTCCTTGCCAGCAGGTACGCCTCACTTGACCCCTACCG<br>TGCAGTGACTCACAATAAGGGAATCATGAATGGCATTGAGGCGGTAGTACTTGCAACAGGGAATGATACCAGAGCTGTTAGTGCTTCTTGTCACGCTTTTGCAGTTAAA<br>GAAGGGCGTTACCAAGGCCTAACTAGTTGGACCCTGGACGGGGAGCAGTTAATTGGTGAAATCAGTGTACCATTAGCTCTGGCAACTGTCGGTGGCGCCACTAAGGT<br>TTTACCTAAGTCACAGGCTGCAGCCGACTTACTTGCTGTAACCGACGCTAAGGAATTATCTAGAGTAGTCGCCGCCGTAGGCTTAGCTCAAAATTTGGCTGCTTTAAGG<br>GCACTGGTGAGTGAAGGAATTCAGAAAGGCCACATGGCCTTACAGGCACGTTCATTGGCGATGACGGTCGGGGCTACCGGCAAGGAAGTAGAGGCAGTGGCTCAGC<br>AGTTGAAAAGACAAAAGACAATGAACCAGGACAGAGCCTTAGCCATCTTAAACGACTTGCGTAAGCAGGGTGGTGGTGGTAGTGGTGGTGGTGGTTCCCTCGAGGAC<br>TACAAAGACGATGACGATAAGtaaATCCTGAGACCgtagcC |
| MvaE_N-term | GgctacGGTCTCAtatgAAAACTGTCGTGATTATCGATGCTTTGAGGACACCAATCGGAAAGTATAAGGGCTCGCTATCTCAGGTGTCAGCCGTAGACTTGGGCACTCATG<br>TTACAACTCAGTTGTTGAAGCGTCACTCAACAATCAGTGAGGAAATAGACCAGGTAATCTTCGGTAATGTACTTCAGGCGGGCAACGGTCAGAACCCCGCTAGGCAGA<br>TCGCTATCAACAGCGGACTATCACACGAAATACCTGCTATGACCGTTAATGAGGTATGTGGCTCAGGAATGAAGGCCGTTATCTTGGCCAAGCAACTGATCCAGTTAG<br>GAGAGGCTGAGGTATTGATTGCTGGTGGTATCGAGAACATGTCACAGGCCCCAAAATTGCAGAGATTCAACTACGAGACAGAGTCATATGACGCTCCCTTTAGCAGTA<br>TGATGTATGACGGGCTTACTGATGCTTTCAGCGGTCAGGCAATGGGCCTAACGGCCGAAAACGTTGCGGAGAAGTATCATGTCACAAGAGAGGAGCAAGACCAATTC<br>TCAGTGCATTCTCAATTGAAGGCCGCACAGGCACAAGCAGAAGGAATCTTTGCAGATGAAATCGCCCCTCTAGAGGTCAGCGGTACACTAGTAGAGAAAGACGAGGG<br>AATTAGGCCCAATAGCTCAGTTGAGAAATTGGGAACACTTAAAACAGTTTTCAAAGAGGATGGCACTGTGACAGCCGGGAATGCCTCAACTATAAATGACGGGGCATC<br>TGCCTTGATAATTGCAAGTCAGGAGTATGCTGAGGCTCACGGGTTGCCCTATCTGGCGATCATCAGGGATTCAGTCGAGGTCGGCATCGACCCAGCATACATGGGTAT<br>CAGTCCAATCAAAGCTATCCAGAAATTATTAGCCCGTAACCAGCTGACAACTGAGGAGATAGACCTGTACGAAATTAACGAGGCATTCGCCGCAACCTCAATAGTAGTA<br>CAGAGAGAATTGGCTCTGCCTGAGGAGAAGGTAAACATTTATGGTGGCGGCATTTCGCTGGGACATGCGATCGGGGCTACTGGTGCCAGGCTTCTGACGAGTTTATC<br>TTACCAACTAAACCAAAAGGAAAAGAAATATGGTGTAGCCAGTCTTTGCATTGGTGGTGGACTTGGTTTAGCTATGTTATTAGAGCGTCCACAGCAGCAGGGTGGTGGT<br>GGTAGTGGTGGTGGTGGTTCCCTCGAGGACTACAAAGACGATGACGATAAGtaaATCCTGAGACCgtagcC |
| MTH1ΔT | GgctacGGTCTCAtATGTTTGTTTCACCACCACCAGCTACTTCTAAAAATCAAGTTTTGCAAAGACGTCCATTGGAATCTACTAATAGTAATCATGGTTTTGCTTCTTCTTTG<br>CAAGCTATTCCAGAAAATACTATGAGTGGCAGTGATAATGCTTCTTTTCAAAGTTTGCCATTGTCAATGTTTTCTGCTCCATCTACTGTTCACACTCAATTGACTAATGAC<br>TCTTCTTTCTCTGAATTTCCTAATCACAAGTTGATCACTAGAGTTTCTCTGGATGAAGCTTTGCCAAAAACTTTTTATGACATGTATTCTCCAGATATTTTGTTGGCTGACC<br>CATCTAATATTTTGTGTAATGGTCGTCCAAAGTTTACAAAGAGAGAGTTGTTGGATTGGGATCTTAATGATATTAGATCTTTGTTGATTGTTGAGAAGTTAAGACCAGAAT<br>GGGGTAATCAATTGCCAGAAGTTATTACTGTTGGTGATAATATGCCACAGTTTAGATTGCAATTGTTGCCATTGTATTCTTCTGATGAGACAATTATCGCTACTTTGGTTC<br>ATTCTGATCTGTACATGGAGGCTAATTTGGATTATGAATTCAAATTGACCTCTGCAAAATATACAGTTGCGACCGCTAGAAAAAGACATGAGCATATTACTGGTAGAAAT<br>GAAGCTGTTATGAATTTGTCTAAACCAGAATGGAGAAATATCATCGAAAATTACTTGTTGAATATTGCTGTTGAGGCTCAATGTAGATTTGATTTCAAACAAAGATGTTCT<br>GAATATAAGAAATGGAAGTTGCAACAATCTAATTTGAAAAGACCAGACATGCCACCACCATCTATTATTCCAAGAAAAAATTCTACAGAAACAAAATCTCTTCTGAAAAAG<br>GCTTTGTTGAAGAATATTCAATTGAAAAATCCAAATAATAATCTTGATGAATTAATGATGAGATCATCTGCTGCTACAAATCAACAAGGTAAAAATAAAGTTTCTTTGTCTA<br>AAGAAGAAAAGGCTACTATTTGGTCACAATGTCAAGCTCAAGTTTACCAAAGATTGGGTTTGGATTGGCAACCAGATTCAGTTTCTTAAATCCTGAGACCgtagcC |

**Supplementary Table 4.** The list of yeast strains used in this study.

| Description | Genotype | Source |
|---|---|---|
| CEN.PK2-1D | MATa ura3-52 leu2-3112 trp1-289 his3Δ1 | EUROSCARF |
| Δ*ald6* | CEN.PK2-1D ald6 ::NatMX6 | This study |
| WT (pRS316) | CEN.PK2-1D (pRS316—empty vector) | This study |
| Δ*ald6* (pRS316) | Δ*ald6* (pRS316—empty vector) | This study |
| WT (pI2M114) | WT (pI2M114—Se-acsL641P, Ef-MvaS, Ef-MvaE_N-term, Ef-MvaE_C-term) | This study |
| Δ*ald6* (pI2M114) | Δ*ald6* (pI2M114—Se-acsL641P, Ef-MvaS, Ef-MvaE_N-term, Ef-MvaE_C-term)) | This study |
| WT (pRS316, pI2M169) | WT (pRS316, pI2M169— MTH1ΔT) | This study |
